# Network-based assessment of the vulnerability of Italian regions to bovine brucellosis

**DOI:** 10.1101/296434

**Authors:** Alexandre Darbon, Eugenio Valdano, Chiara Poletto, Armando Giovannini, Lara Savini, Luca Candeloro, Vittoria Colizza

**Affiliations:** INSERM, Sorbonne Université, Institut Pierre Louis d’Epidémiologie et de Santé Publique IPLESP, F75012 Paris, France; Departament d’Enginyeria Informàtica i Matemàtiques, Universitat Rovira i Virgili, Tarragona, 43007, Spain; Istituto Zooprofilattico Sperimentale dell’Abruzzo e del Molise, Teramo, Italy

**Keywords:** Brucellosis, networks, animal trade, vulnerability, epidemic model

## Abstract

The endemic circulation of bovine brucellosis in cattle herds has a markedly negative impact on economy, due to decreased fertility, increased abortion rates, reduced milk and meat production. It also poses a direct threat to human health. In Italy, despite the long lasting efforts and the considerable economic investment, complete eradication of this disease still eludes the southern regions, as opposed to the northern regions that are disease-free. Here we introduced a novel quantitative network-based approach able to fully exploit the highly resolved databases of cattle trade movements and outbreak reports to yield estimates of the vulnerability of a cattle market to brucellosis. Tested on the affected regions, the introduced vulnerability indicator was shown to be accurate in predicting the number of bovine brucellosis outbreaks, thus confirming the suitability of our tool for epidemic risk assessment. We evaluated the dependence of regional vulnerability to brucellosis on a set of factors including premises spatial distribution, trading patterns, farming practices, herd market value, compliance to outbreak regulations, and exploring different epidemiological conditions. Animal trade movements were identified as a major route for brucellosis spread between farms, with an additional potential risk attributed to the use of shared pastures. By comparing the vulnerability of disease-free regions in the north to affected regions in the south, we found that more intense trade and higher market value of the cattle sector in the north, likely inducing more efficient biosafety measures, together with poor compliance to trade restrictions following outbreaks in the south were key factors explaining the diverse success in eradicating brucellosis. Our modeling scheme is both synthetic and effective in gauging regional vulnerability to brucellosis persistence. Its general formulation makes it adaptable to other diseases and host species, providing a useful tool for veterinary epidemiology and policy assessment.

## Introduction

Brucellosis is the most common zoonotic disease worldwide, severely hindering livestock productivity and human health (Corbel, 1997; Pappas et al., 2006; Ariza et al., 2007; Franc et al., 2018). Bovines are the natural host for *Brucella abortus*, the pathogen causing bovine brucellosis, a highly contagious sub-acute and chronic disease leading to abortion, premature births, weak offspring, retained placenta, or infertility (Corbel, M.J., 2006). Transmission between animals generally occurs by direct contact and environmental contamination following an abortion. The disease is the cause of severe economic losses as a consequence of late term abortion, substantial decline in milk and meat production, reduced fertility, and the induced effects of control strategies such as the loss of infected animals to culling and restrictions in commerce.

Despite the great progress in controlling and eradicating the disease in many areas of the world (Pappas 2006), there still remain regions where the infection is endemic in livestock. In Europe the vast majority of Member States (19 out of 28, 68%) are currently declared officially free from bovine brucellosis (OBF) (EFSA and ECDPC, 2017), following successful eradication campaigns co-financed by the European Union. The disease still persists in the Southern countries close to the Mediterranean basin where it is responsible for the largest notification of infections in humans in Europe, often leading to severe cases requiring hospitalizations. Eradication is dramatically hampered by several factors: the sub-acute nature of the disease that leads to an initial phase following the infection that is often not apparent (silent spreading phase); the lack of specificity of serological tests that may cross-react with other bacteria (Fadeel et al., 2011; Varshochi et al., 2011); the performance of control activities and the lack of full compliance to reporting and regulations in some contexts (Calistri et al., 2013; Graziani et al., 2013; Bronner et al., 2014).

Italy currently has OBF status for 11 regions and 9 provinces, localized in the center and north of the country (EFSA and ECDPC, 2017). The first eradication strategy was introduced in 1994 through a strict test-and-slaughter policy when the disease was endemic in the entire country (Italian Ministry of Health, 1994), after the discontinuation of animal vaccination. Efficient progress towards eradication was made in the regions of the north and center that progressively gained the OBF status. Despite the continuous and considerable efforts of the veterinary services and the implementation of stricter regulatory policies in 2006 (Italian Ministry of Health, 2006), the southern part still remains affected by the disease with a considerable persistence in the regions of Campania, Calabria, Puglia, and Sicilia (Figure 1) (EFSA and ECDPC, 2017).

**Figure 1.**
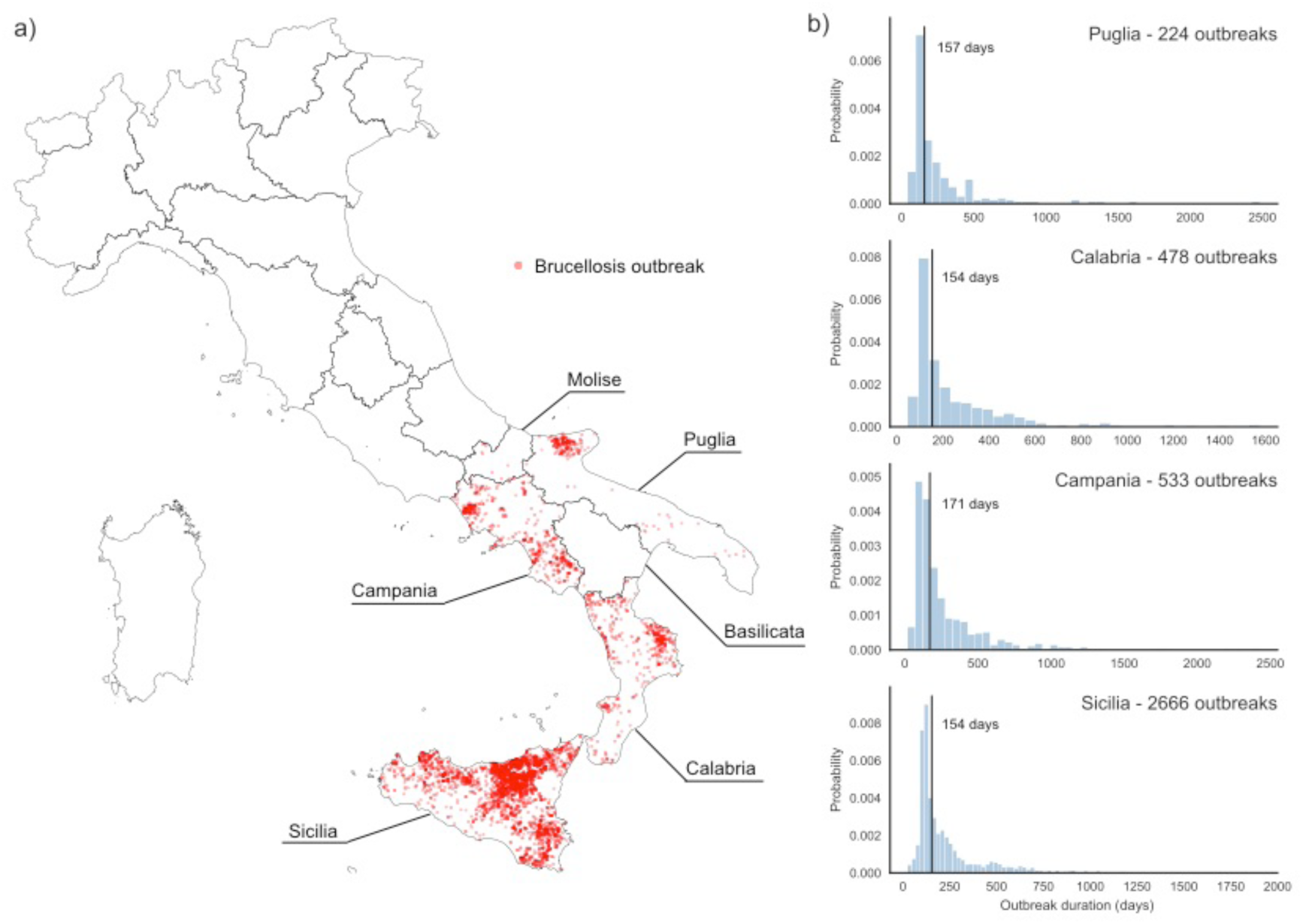
Bovine brucellosis outbreaks in Italy. a) Location of bovine brucellosis outbreaks in Italy in the period February 1983 - September 2015. Only the six most southern regions experienced outbreaks in the period under study. b) Distribution of outbreak duration per region, in the four most affected regions. Solid lines show median values.

Previous studies identified important drivers for brucellosis persistence in the affected countries in Europe. Contiguous spread from previous outbreaks was generally identified as the main source of new infections (Godfroid and Käsbohrer, 2002). Pathways for brucellosis spatial transmission in Spain and Northern Ireland were contacts between herds, in particular from pasture rental practices, neighboring herds, or through cattle movements between farms (Abernethy et al., 2006; Stringer et al., 2008; Dias et al., 2009; Abernethy et al., 2011; Cowie et al., 2014). The only study on risk factors for bovine brucellosis in Italy was conducted in the insular region of Sicilia and showed that herd size and beef or mixed production types were associated with higher risk of persistence of the disease in the region (Calistri et al., 2013). The number of animals introduced from other herds instead did not seem to play an important role.

The aim of this study was to quantitatively assess the vulnerability of Italian regions to bovine brucellosis to explain the extremely diverse epidemiological situation reported in the country, with strong regional differences. Adopting a network perspective for veterinary epidemiology and policy assessment (Dubé et al., 2009; Martínez-López et al., 2009), we developed a mathematical model describing the spread of bovine brucellosis from farm to farm associated with the movement of infected animals in the region. The model is based on the complete database of daily cattle trade movements for the whole country from January 2006 to December 2012, and it is parameterized with outbreak data reported to the Italian National Veterinary authority in the period 1983-2015. We introduced a measure of vulnerability quantifying the risk for brucellosis persistence in the affected regions, and compared OBF areas with non-OBF areas accounting for premises spatial distribution, trading patterns, farming practices, outbreak status, herd market value, and exploring different epidemic conditions.

## Material and Methods

### Cattle trade movement data

Data on cattle trade movements were obtained from the Italian National Database for Animal Identification and Registration (NDB) that is managed by the Istituto Zooprofilattico Sperimentale dell’Abruzzo e del Molise (IZSAM) on behalf of the Italian Health Ministry (“Sistema Informativo Veterinario,” n.d.). The database details the movements of the entire Italian population of bovines among animal holdings, providing a comprehensive picture of where cattle have been kept and moved within the country. Each movement record reports the unique identifier of the animal, the identifiers of the holdings of origin and destination, and the date of the movement. Holdings are georeferenced.

Here we examined the daily records from January 1, 2006 to December 31, 2012. A total of 11,775,842 bovines were tracked, counting for 3,693,593 recorded displacements among 190,596 animal holdings in Italy (Table 1).

**Table 1.** Basic description of regional cattle trade systems in Italy.

| Region |  | #premises | #trading links |  | #moved bovines |  | in-degree | in-strength |
| --- | --- | --- | --- | --- | --- | --- | --- | --- |
|  |  |  | *total* | *avg/y* | *total* | *avg/y* | *avg/y* | *avg/y* |
| NORTH | Piemonte | 18,993 | 598,155 | 85,450 | 1,669,615 | 238,516 | 3.99 | 16.87 |
|  | Valle d'Aosta | 1,632 | 48,876 | 6,982 | 103,144 | 14,735 | 3.20 | 9.10 |
|  | Lombardia | 22,891 | 759,936 | 108,562 | 2,929,995 | 418,571 | 3.35 | 23.44 |
|  | Trento Alto-Adige | 11,637 | 400,109 | 57,158 | 608,486 | 86,927 | 2.31 | 4.25 |
|  | Veneto | 19,492 | 506,377 | 72,339 | 2,173,769 | 310,538 | 2.32 | 17.44 |
|  | Friuli Venezia Giulia | 3,133 | 108,662 | 15,523 | 221,740 | 31,677 | 1.58 | 6.40 |
|  | Liguria | 1,834 | 13,359 | 1,908 | 28,091 | 4,013 | 1.03 | 2.18 |
|  | Emilia Romagna | 10,795 | 532,423 | 76,060 | 1,622,371 | 231,767 | 2.10 | 16.16 |
| CENTER | Toscana | 6,075 | 51,586 | 7,369 | 146,927 | 20,990 | 1.39 | 4.68 |
|  | Umbria | 4,862 | 37,460 | 5,351 | 98,684 | 14,098 | 1.26 | 3.94 |
|  | Marche | 5,995 | 49,346 | 7,049 | 123,476 | 17,639 | 1.30 | 4.00 |
|  | Lazio | 15,098 | 124,222 | 17,746 | 354,362 | 50,623 | 1.30 | 4.47 |
|  | Abruzzo | 7,021 | 52,625 | 7,518 | 107,585 | 15,369 | 1.39 | 3.36 |
|  | Sardegna | 10,642 | 66,542 | 9,506 | 410,764 | 58,681 | 1.19 | 8.02 |
| SOUTH | Molise | 3,821 | 29,528 | 4,218 | 55,093 | 7,870 | 1.18 | 2.46 |
|  | Campania | 14,992 | 92,117 | 13,160 | 214,501 | 30,643 | 1.32 | 3.56 |
|  | Puglia | 5,533 | 58,335 | 8,334 | 191,759 | 27,394 | 1.39 | 6.83 |
|  | Basilicata | 3,729 | 27,697 | 3,957 | 106,833 | 15,262 | 1.10 | 4.38 |
|  | Calabria | 9,370 | 36,545 | 5,221 | 128,280 | 18,326 | 1.13 | 4.52 |
|  | Sicilia | 13,051 | 99,693 | 142,412 | 480,367 | 68,624 | 1.60 | 9.10 |

### Network-based spatial transmission model

Cattle movements were represented in terms of a network with nodes corresponding to premises, and directed edges representing animal movements between two premises (Dubé et al., 2009; Martínez-López et al., 2009). The static projections of Italian network aggregated over a year were provided by (Natale et al., 2009), and its dynamical properties considering multiple timescales were assessed in (Bajardi et al., 2011; Valdano et al., 2015c).

Given the fundamental importance of causality in the sequence of trading links for the transmission of cattle diseases (Bajardi et al., 2011; Vernon and Keeling, 2009), we built the full time-resolved directed weighted network based on daily movement records. We denoted with *W*_*t*_ the weighted adjacency matrix representing the network at the daily snapshot *t* in the time period under study. The matrix element *W*_*t,ij*_ corresponds to the number of animals moved from holding *i* to holding *j* at time *t*, and it is equal to zero if no animals are moved.

Since we are interested in the transmission and persistence of brucellosis at the regional level, rather than the detailed dynamics within a single farm, we modeled spatial transmission from farm to farm considering animal holdings as discrete epidemiological units. The spread on the dynamical network was modeled using a simple susceptible-infectious-susceptible (SIS) approach for disease progression (Keeling and Rohani, 2007) where premises can be in one of the following two states: susceptible (S), if the farm is brucellosis-free, and infectious (I), if the farm experiences a brucellosis outbreak. Transmission may occur from an infected farm *i* to a susceptible one *j* upon trading animals at time *t* with a probability *Λ*_*t,ij*_ = 1 − (1 − λ)^*W*_*t,ij*_^, where *W*_*t,ij*_ counts the number of animals moved and *λ* indicates the per-animal risk of transmission. The expressed probability *Λ*_*t,ij*_ thus accounts for an increased risk associated to a larger number of traded animals (Bajardi et al., 2011; Valdano et al., 2015b). An infected farm can clear the infection with rate *μ* whose inverse measures the average duration of a brucellosis outbreak in a farm. Once disease-free, the farm recovers the S status and can be infected again with brucellosis, thus experiencing successive outbreaks.

To assess the spatial heterogeneity of the brucellosis epidemiological situation in Italy at the regional level, we considered regional sub-networks of the national cattle trade network, each of them including all premises and all trading links within the region (Table 1).

We also measured basic network indicators for each animal holding: the in-degree of farm *i* counting the number of farms which farm *i* receives cattle from (and similarly for the out-degree counting outgoing links); the in-strength of farm *i* counting the number of incoming animals in the farm (and similarly for the out-strength counting the number of animals moved out of the farm). These quantities can be defined at a given time resolution (e.g. during a day, a week, a month, etc.) (Bajardi et al., 2011), and here we considered the yearly timescale. We then computed averages and standard deviations for these indicators over the years of the dataset under study, and over all farms in the region (Table 1).

### Model parameterization

Data on brucellosis control activities were collected from the Italian National Animal Health Managing Information System, an informatics system developed by IZSAM on behalf of the Ministry of Health to monitor the progress of the eradication plan and measure the degree of achievement of its objectives (Italian Ministry of Health, 2012; “Sistema Informativo Veterinario,” n.d.). The database reports on all surveillance and eradication activities and their results at farm and individual animal level. We used records of bovine brucellosis outbreaks occurring in Italy between February 1983 and September 2015. For each outbreak, we considered data reporting the identifiers of the affected animal holding, its production type, and the dates of start and end of the outbreak, based on the date of first infection detection and date of clearing of the infection, respectively.

To account for different regional contexts, we parameterized the transmission model in the brucellosis-affected regions by setting the recovery rate *μ* equal to the inverse of the median outbreak duration reported for each region, and listed in Table 2. We also explored fluctuations of this value within the 95% confidence intervals.

**Table 2.** Outbreaks and their duration per region (median and 95% confidence interval).

| **Region** | **# outbreaks** | **Outbreak duration (days)** |
| --- | --- | --- |
| Sicilia | 2666 | 155 [68; 1125] |
| Campania | 533 | 171 [67; 1007] |
| Calabria | 478 | 154 [81; 928] |
| Puglia | 224 | 157 [66; 1039] |
| Molise | 6 | 201 [94; 212] |
| Basilicata | 3 | 104 [61; 115] |
| OBF regions | 0 | 157* |

\*assumed value obtained as average of non-OBF outbreak durations

Since regions in the center and north of Italy have been OBF for many years, no empirical data exist on the duration of the outbreak. We then parameterized the SIS model in those regions using the average of the outbreak durations reported in brucellosis-affected regions. The underlying hypothesis is the comparison of the risk of brucellosis persistence across all regions of Italy under the same epidemiological and veterinary conditions (here measured by the outbreak duration in the southern affected regions).

In addition, we explored a full range of values of the outbreak duration, from 10 days to 10,000 days, the upper limit used for theoretically assessing the trend of vulnerability to brucellosis for varying outbreak durations.

The per-animal transmission rate *λ* is left as a variable to identify the critical conditions for brucellosis persistence in each region, as explained in the following subsection.

### Epidemic vulnerability assessment, control strategies, and compliance

We introduced a measure of vulnerability *v* quantifying the risk for brucellosis persistence in a given region, once the disease is introduced in the region. It is defined as the inverse of the critical value *λ*_*c*_ of the transmissibility *λ, v* = 1*λ*_*c*_, where *λ*_*c*_ quantifies the critical condition for the endemic state of the disease (Anderson and May, 1992; Keeling and Rohani, 2007). If the transmissibility is larger than the critical value, *λ > λ*_*c*_, the model predicts the disease to spread through the cattle trade system and become persistent; if the transmissibility is lower than this value, *λ < λ*_*c*_, the model predicts the disease to go extinct.

To compute the epidemic vulnerability of each region to bovine brucellosis and how it depends on cattle trade patterns and outbreak conditions, we consider the *infection propagator approach* explicitly integrating time-resolved animal movements among premises (Valdano et al., 2015a). The approach is based on the SIS spatial transmission model described above and parameterized with the outbreak data. Adopting a Markov chain perspective (Wang et al., 2003; Gomez et al., 2010) under the assumption of locally treelike networks (Gleeson et al., 2012; Radicchi and Castellano, 2016), the equations describing the SIS propagation among farms connected by time-resolved trading links are:

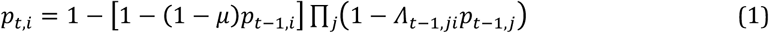

where *p*_*t i*_ is the probability that farm *i* is infected at time *t* and *A*_*t* − 1,*ji*_ is the transmission probability from *j* to *i* at time *t −* 1, defined previously. By mapping the process into a multilayer formalism accounting for both spreading and network dynamics (Valdano et al., 2015a), the infection propagator approach allows for the analytical solution of the linearized Markov chain description of the spreading process, with the epidemic threshold *λ*_*c*_ expressed as the solution of the equation:

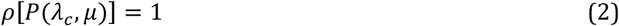

where *ρ* is the spectral radius of the following matrix:

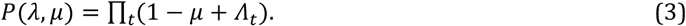

The generic element *P*_*ij*_ represents the probability that brucellosis can spread from an infected farm *i* at the beginning of the time period under study to farm *j* at the end of this period through time-respecting transmission paths along the temporal trading links of the network. For this reason it is denoted as infection propagator. More details on the analytical treatment are provided in (Valdano et al., 2015a, 2015b).

For each region, we compute the brucellosis vulnerability *v* = 1/*λ*_*c*_, with *λ*_*c*_ obtained from Eq. (2) where the infection propagator *P* is parameterized with cattle trade data and brucellosis clearing rate *μ* obtained from regional outbreak data. We compute fluctuations of the vulnerability by taking into account the empirically obtained fluctuations of the outbreak duration per farm (Table 2).

The SIS transmission scenario implicitly considers the implementation of control strategies, as the model allows for a farm to clear the infection after an outbreak. Their specific details, such as e.g. the number and timing of testing and culling of infected animals, cannot be explicitly considered here because of the farm-level description of our transmission model. The promptness of their implementation and their efficacy are however effectively integrated and represented by the duration of the outbreak.

In addition, we measured the compliance to animal movements ban by identifying the infected farms that continue to trade animals while infected and characterize their trading behavior with respect to the same seasonal period in a previous year when they were brucellosis free (also accounting for yearly variations of overall trading volume).

### Statistical analyses

Spearman correlation coefficients were computed to test the correlation between the regional brucellosis vulnerability and a set of different features: (i) the number of brucellosis outbreaks experienced in the region; (ii) animal trade features, such as the average farm in-degree and average farm in-strength; (iii) average regional cattle market value (2016 data) (RICA, -); (iv) farming practices, considering the number of premises with pastures per region (2010 data, one-tailed test, as a decrease in risk induced by pastures is not supported by biological plausibility) (ISTAT, 2011).

In addition, we analyzed the relation between the number of outbreaks experienced by a single farm and the number of herds purchased from infected farms.

## Results

A total of 3,910 outbreaks affecting 3,214 premises were recorded in the period under study in the regions of Puglia, Calabria, Campania, Basilicata, Molise, and Sicilia (Figure 1), with the largest fraction occurring in Sicilia (2,666, 68%). Outbreak duration in a farm was heterogeneously distributed (except for Basilicata and Molise experiencing very few outbreaks), with median values ranging from 104 days in Basilicata to 201 days in Molise (Figure 1 and Table 2), and overall average of 157 days.

A widely heterogeneous vulnerability situation was predicted for brucellosis-affected regions in the south of Italy (Figure 2). Sicilia is by large the region with the highest vulnerability to brucellosis (*v* = 15.9 [4.3; 190.8]), followed by Campania (*v* = 8.2 [2.2; 318.0]). Most importantly, the estimated vulnerability per region was strongly associated with the number of outbreaks recorded in the region (Spearman *r* =0.82, *p* =0.04).

**Figure 2.**
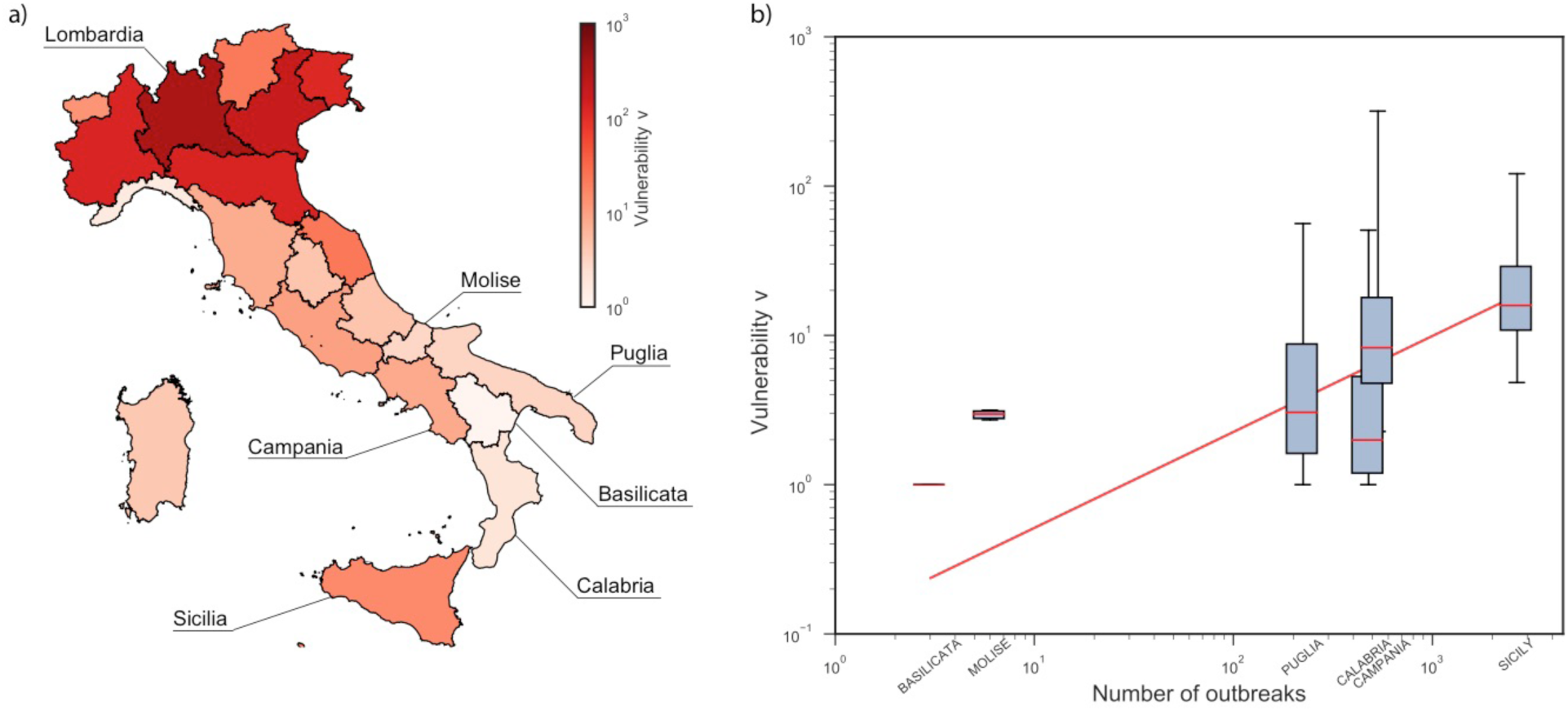
Vulnerability of Italian regions to bovine brucellosis and comparison with number of outbreaks. a) Map of predicted vulnerability per region. The color scale refers to the estimated value of *v* per region, when the model is parameterized with the data of Table 2. b) Relation between vulnerability and number of outbreaks per region in the south. Whisker boxes (red line at the median, box at 50% confidence interval, whiskers at 95% confidence intervals) represent the variation in the predicted vulnerability due to fluctuations in the estimated outbreak duration from data. The red straight line is a log-log fit shown as a guide to the eye.

Risk assessment results were highly heterogeneous also for the regions in the center and north of the country, with vulnerability values ranging from *v* = 1.6 in Liguria to *v* = 364.8 in Lombardia, the region with the highest predicted risk in the country. Unexpectedly, we found considerably higher vulnerability estimates in the northern OBF regions compared to the affected regions in the south, once the model was parameterized in those regions with outbreak statistics from the south.

A large difference in trading and farming presence was reported between the north and the rest of the country. Almost half of the total number of premises (90,407, 47%) was located in the northern regions, and the vast majority of animal movements originated in those same regions (2,967,897, 80%, Figure 3). On the other hand, the proportion of animals traded within each region compared to across-region movements was quite stable across the different areas.

**Figure 3.**
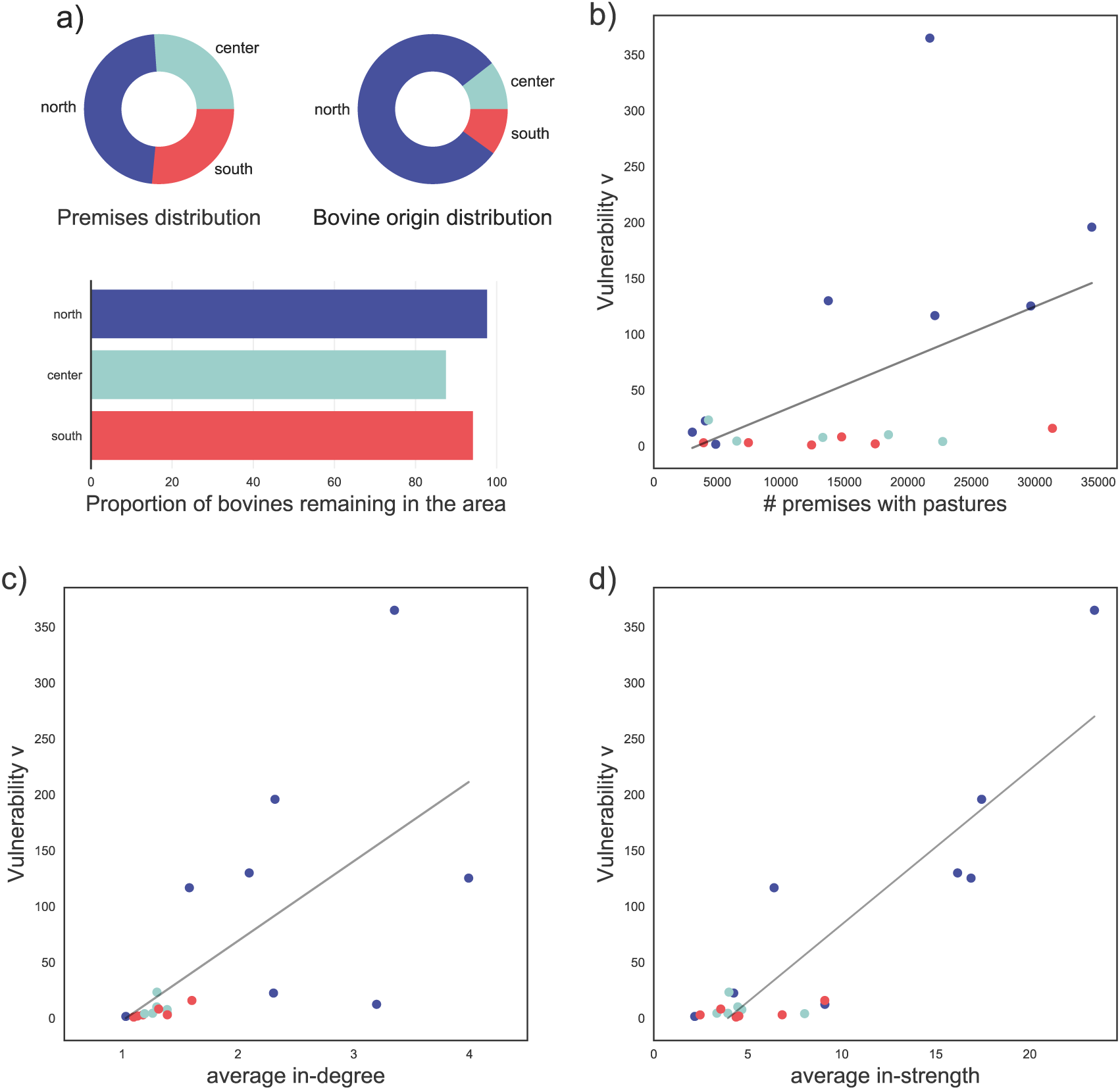
Impact of trading and farming practices on regional vulnerability to brucellosis. a) Distribution across the country (north, center, and south) of: number of premises, number of bovines moved out of a farm, proportion of animals traded within the area. b), c), d) Relation between vulnerability and: number of premises with pastures (panel b), average in-degree (panel c), average in-strength (panel d) per region. Each dot represents a region and is color-coded according to its geographic position (north, center, south, as in plots of panel a). Black straight lines are linear fits shown as guides to the eye.

The regional vulnerability to brucellosis had a strong association with the average in-degree *(r* = 0.85, *p* < 10^−5^, panel c of Fig. 3) and the average in-strength of a farm in the region *(r* = 0.65, *p* < 10^−2^, panel d). The association was moderate with the number of premises with pastures in the region *(r* = 0.4, *p* = 0.04, panel b of Fig. 3).

Given that no accurate parameterization is possible for the transmission model in the OBF regions because of the absence of outbreaks, we synthetically explored outbreak durations ranging from 1 to 10^4^ days. Vulnerability was found to increase with the average outbreak duration, as expected (Figure 4). Lombardia was at a consistently higher risk (two orders of magnitude) compared to Sicilia. We found that the vulnerability estimated in Sicilia based on the region’s brucellosis outbreak duration of 157 days would be recovered in Lombardia if outbreaks lasted on average 12 days.

**Figure 4.**
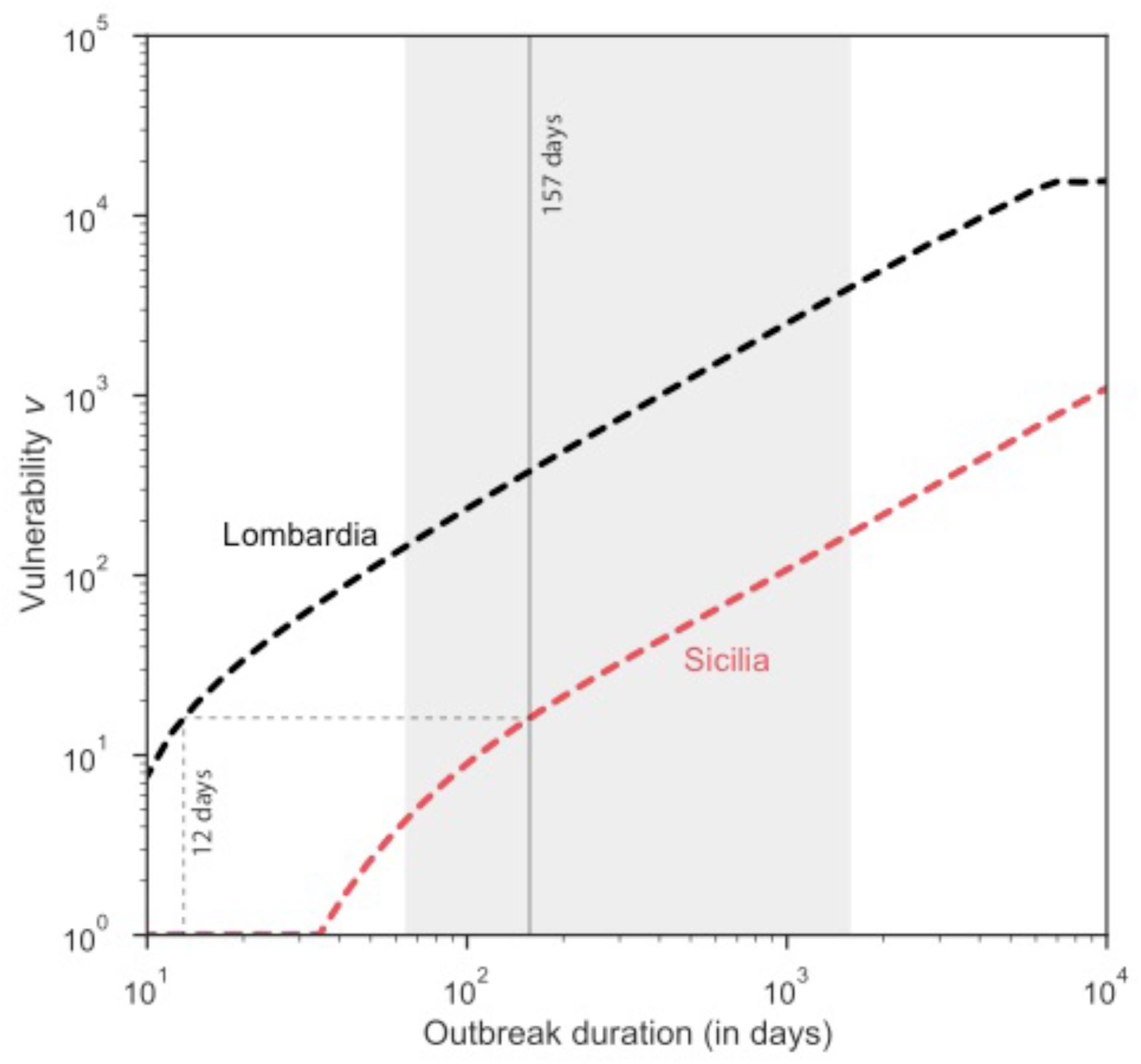
Impact of outbreak duration on regional vulnerability to brucellosis. Relation between vulnerability and outbreak duration for Sicilia (non-OBF region at highest predicted risk) and Lombardia (OBF region at highest predicted risk). Median duration of brucellosis outbreak in the south (vertical grey line) is shown together with its 95% confidence interval (vertical grey area).

Regional vulnerability to brucellosis (parameterized on southern outbreaks) was moderately associated with the average regional cattle market value *(r* = 0.56, *p* = 0.01, Figure 5), indicating that the most vulnerable regions are also the highest capitalized markets.

**Figure 5.**
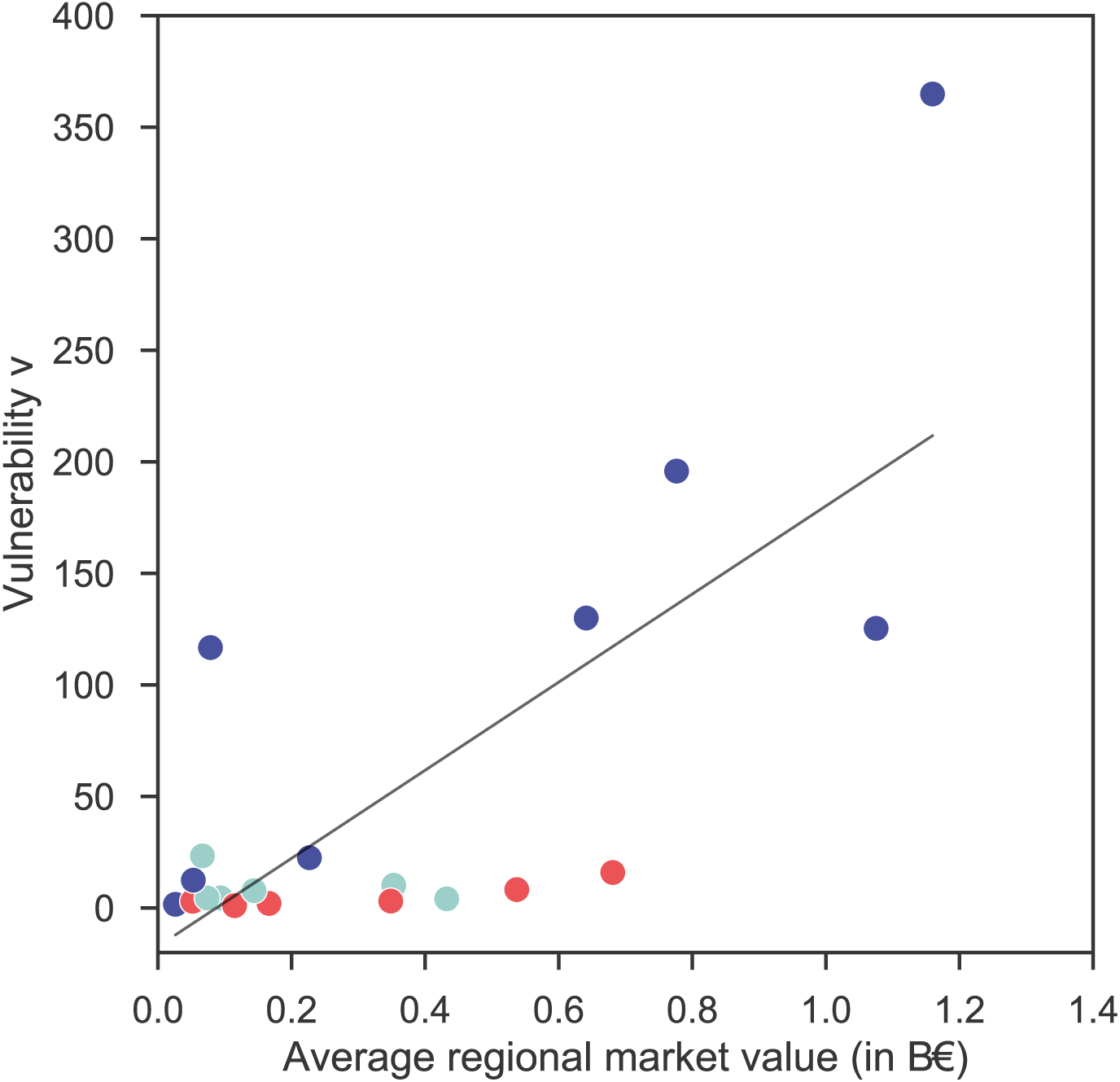
Impact of cattle market value on regional vulnerability to brucellosis. Relation between vulnerability and average cattle market value per region. Each dot represents a region and is color-coded according to its geographic position (north, center, south, as in Figure 3). The black straight line is a linear fit shown as a guide to the eye.

Finally, we analyzed the compliance of farmers to the regulatory framework put in place in case of brucellosis outbreak declared in affected premises. Despite the imposed movement ban following the detection of the infection, we found that 15% of the infected premises continued to trade (Figure 6). Approximately half of them traded at least 50% or more of the animals compared to the same period of the year while non-infected, with 17% increasing their trade by 150%. The number of outbreaks occurring in a given farm was strongly associated with the number of traded herds originated from infected premises *(r* = 0.88, *p* < 10^−5^).

**Figure 6.**
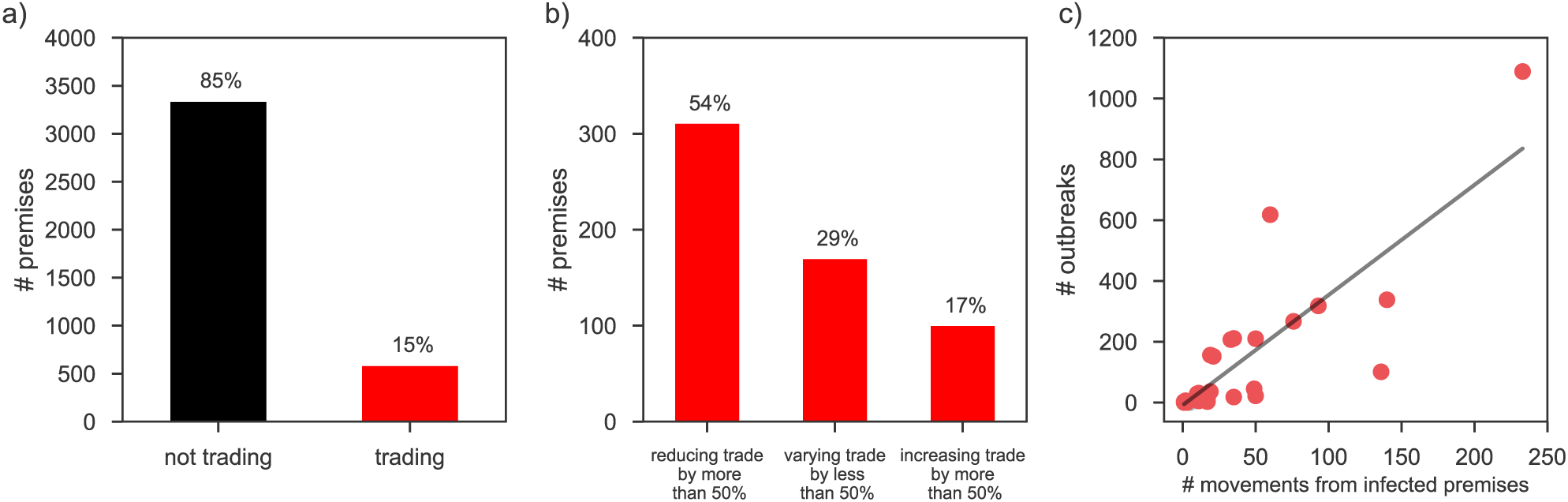
Compliance to trade bans and impact on bovine brucellosis diffusion. a) Number of infected premises according to their trading behavior while infected. b) Cattle movement volume variation for infected premises that trade when infected. c) Relation between number of outbreaks experienced by farms in a given province and number of movements received from infected premises. The black straight line is a linear fit shown as a guide to the eye.

## Discussion

Despite continuous progress in the control of bovine brucellosis in Italy, leading to a decline in prevalence from 0.016 to 0.08 in the period 2009-2016 (“Sistema Informativo Veterinario,” n.d.), complete eradication has not been achieved and 21% of provinces in the country are still affected, with a significant economic damage to the agricultural and food industries. Here we introduced a novel mathematical approach to estimate the vulnerability of regional cattle populations to brucellosis persistence, based on the spatial transmission of the disease through animal movements. Parameterized on empirical data from the affected regions, our approach is able to capture the large epidemiological heterogeneity reported in these regions and independently explain the number of brucellosis outbreaks (re-)occurring in the south of Italy. This reveals that the introduced vulnerability is an accurate indicator of the likelihood of disease persistence in a given area, showing the potential of a synthetic assessment for veterinary epidemiology based on animal movements and disease spatial transmission.

Comparing the affected regions with OBF ones through the vulnerability indicator becomes instead problematic as it suffers from an incomplete parameterization. Having eradicated the disease since many years, there is indeed no outbreak data to accurately inform our transmission model and specifically Eq. (3). Our findings showed that, in presence of endemic circulation of brucellosis in the whole country with infection clearing rates in the farms similar to what recorded in the south, regions of the north would be considerably more vulnerable than those in the south. Several factors drive this result. Distribution of farms is denser in the north, an area of the country that is highly industrialized and characterized by intensive farming and production. Even though the proportion of animals moved within the region is similar throughout the country, trading is more frequent in the north where premises exchange animals with a larger number of holdings and in larger batches compared to center and south. The number of connections and the total volume of traffic going through a node are well-established drivers for infection spread in a host population (Anderson and May, 1992; Pastor-Satorras and Vespignani, 2001; Lloyd and May, 2001; Moreno et al., 2002; Keeling and Rohani, 2007; Barrat et al., 2008; Dubé et al., 2009; Martínez-López et al., 2009; Vernon and Keeling, 2009). Farms that are on average better connected also increase their possibility to contract the infection from neighboring farms, and, on their turn, they can transmit the disease to a larger portion of the trading system, strongly increasing its vulnerability. In addition, northern regions did not suffer from travel bans reducing commercial trade in the studied period because of their OBF status, compared to southern regions. These results clearly show the role of animal movements in contributing significantly to the maintenance of the disease, confirming previous results obtained in Spain identifying the number of purchased animals as a risk factor for brucellosis persistence (Cowie et al., 2014; Dias et al., 2009).

Other types of animal movements beyond trade may play an important role in the disease diffusion, such as movements to pastures (Dias et al., 2009; Abernethy et al., 2011; Calistri et al., 2013; Cowie et al., 2014). These movements are not included in the cattle trade database used in this study as Italian regulations do not require their compulsory tracking. Using a different dataset, we were able however to find a moderate association of the brucellosis vulnerability of each region with the number of premises with pastures. Direct contact between cattle at pasture was found to be the most likely driver for bovine brucellosis transmission in Northern Ireland (Abernethy et al., 2011), where several neighboring herds reported epidemics breaking out during summer (when they more likely moved to pastures), and epidemiological investigations identified deficiency boundary fencing in the area. Similar results were obtained also in Spain, where the number of farmers sharing summer pastures was identified as a significant risk factor for brucellosis (Dias et al., 2009; Cowie et al., 2014). In the Italian region of Sicilia, a previous study indirectly concluded that the habit of moving animals to shared pastures may underlie the observed clustering of infected herds in specific locations, without however providing a statistical analysis for lack of data (Calistri 2013). Here we show that the possible risk provided by the specific type of husbandry adopted involving the use of pastures is aligned with the vulnerability computed on commercial trade, thus strengthening our overall risk assessment. The region with the largest number of premises with access to pastures is indeed Sicilia with more than 31,000 premises (ISTAT, 2011), followed by regions in the north such as Piemonte and Veneto also having indeed a high vulnerability. Though eradicating a disease in islands like Sicilia is expected to be eased by the possibility of better isolating the territory and defend it from external importation, these areas are also the ones characterized by a breakdown of the use of the agricultural territory that facilitates uncontrolled contact between neighboring herds: for example, more than 40% of the farming territory is used for pastures in Sicilia, compared to only 18% in the center regions (ISTAT, 2016).

These findings analyzed the vulnerability of Italian regions under the same epidemiological and control conditions throughout the country, i.e. a hypothetical and non-realistic scenario for OBF regions. The synthetic exploration of the average brucellosis outbreak duration in a farm however showed that Lombardia, the highest vulnerability northern OBF region, would experience a risk similar to Sicilia for infections lasting for about one tenth of the average duration in a Sicilian farm, i.e. approximately 10 days. Given that brucellosis has been eradicated in Lombardia since many years, these findings seem to point to a considerably prompter and more efficient control in the regions of the north and center than those in the south, thus minimizing the duration of an outbreak following an importation and effectively preventing its further spread. Since farmers are required to declare all clinical signs associated to brucellosis according to national regulations, the duration of the outbreak considered in this study is not only a measure of disease progression in time in the affected herd, but also quantifies the response of the farmer and of veterinary services called for its control, having an inevitable impact on brucellosis persistence. In Sicilia, Calistri *et al.* found that the time between detection of the infection and slaughtering of infected animals influences the health status of the herd in the following testing (Calistri et al., 2013).

Measuring promptness and efficacy of interventions directly, for instance through a survey, might provide additional evidence to further support our findings. Very few studies have however considered in detail the possible variety of biosecurity practices adopted on premises to reduce transmission between farms through direct contacts (e.g. investigations or testing prior to animal purchases), indirect contacts (e.g. reducing possible contamination of equipment, personnel, vehicles), or transmission within farms (e.g. enhanced cleaning, hygiene, change of material, etc.). This is mainly due to the large costs and efforts required to conduct biosecurity surveys to gather information on preventive actions at the premises level and to farmers’ acceptability. Few published data exist for the UK (Davison et al., 2003; Brennan and Christley, 2012), the US (Brandt et al., 2008; Hoe and Ruegg, 2006), and Sweden (Nöremark et al., 2010). Little is available for cattle farms in Italy and is generally focused on specific events or activities (Fratini et al., 2010), or on affected regions (Graziani et al., 2013) and thus not generalizable. To overcome the lack of biosecurity data, we thus tested the correlation between vulnerability and regional market value of the cattle compartment, finding a moderate association. This indirectly suggests that premises at a considerably higher risk because of their intensive farming practices are also the ones that implement stricter, prompter and more efficient preventive and control policies to protect the large capital invested in their production. Circulation of brucellosis on northern premises would cause dramatically larger economic losses compared to extensive farming in the south.

Efficacy of interventions is also related to the compliance of farmers to regulations. This is generally outlined in official reports monitoring the progress of control activities (Graziani et al., 2013), through for example measures of the percentage of effectively tested herds on the total subject to control, or of the percentage of infected animals that are slaughtered. An important part of brucellosis control is the ban on movements as soon as an infection is detected in a given holding. By crossing the database of animal movements (NDB) and the database of brucellosis surveillance and eradication activities (SANAN) (“Sistema Informativo Veterinario,” n.d.), our study allowed for the first time the assessment of farmers’ compliance to restrictions on trade while experiencing an ongoing outbreak. Approximately 15% of farmers with infected herds continued to trade despite the policy in place, and half of them moved about the same number of animals or more than they were used to move in the same seasonal period. Most importantly, the number of outbreaks occurring in a given premises is clearly associated with the number of incoming traded herds received from infected premises. Such risk was previously assessed only in Northern Ireland, where it was found to be 19 times higher compared with animals from herds with no history of infection (Abernethy et al., 2011). Our study confirms that movements are therefore significant for disease dissemination and persistence. This was not previously observed in the analysis of brucellosis risk factors in Sicilia (Calistri et al., 2013), likely because of confounding factors such as the large number of pastures present in the sole region under study.

The strong reluctance of producers in complying with eradication programs is expected to be related to the large losses incurred. The total cost for brucellosis eradication in the region of Lazio in central Italy from 2007 to 2011 was estimated to be 6M Euros with 6.5% contribution of the European Union (Caminiti et al., 2016), for an average number of 236,262 cattle heads. While including culling, representing 0.5% of the overall figure, this estimate did not however consider costs associated with movement bans or additional losses for farmers such as decreased milk and meat production. It is important to note that gaining OBF certification halved the overall cost, thus showing the importance of attaining disease eradication through a collective and proactive approach, for both national authorities and individual producers. In addition to measures to improve compliance, different management strategies can be envisioned, some of them already adopted by other countries, such as e.g. testing prior to movements, quarantine of animals following trade, or cattle restrictions on larger areas, i.e. extending travel bans from the farm level to the province or regional area. This is especially important for diseases characterized by a silent spreading phase with a latent sub-clinical infection that may hide epidemiological links between premises due to the long interval between infection and disclosure.

Finally, we stress the importance of cross-matching different databases such as the one for bovine identification that tracks animal movements and the database for the continuous monitoring of surveillance and control activities, in order to uncover unexpected behaviors weakening the progress of efficient brucellosis eradication, as we showed here. We suggest that such cross-matching is performed routinely by veterinary authorities to considerably enrich the knowledge on the health status of animal populations and improve the accuracy of the evaluation of veterinary policies for updates and replanning.

This study is affected by some limitations. First, we did not consider interactions between cattle and wildlife that may act as a reservoir for Brucella pathogens. Spillovers from wildlife are known to pose a greater risk for domestic pigs than for cattle, due to the presence of hares and wild boars infected with *Brucella suis* (Godfroid and Käsbohrer, 2002). In Spain, wild ruminants were not identified as a significant brucellosis reservoir for livestock (Munoz et al., 2010). Contacts with wildlife also strongly depend on farming practices. Indoor farming is the most common breeding practice adopted in Italy with the effect of strongly limiting brucellosis transmission through interactions with wildlife.

Second, our assessment of the vulnerability of cattle population to brucellosis refers to the risk of persistence within the country, once the disease has already been introduced. Thus, it does not account for the probability of importation that may vary throughout the country. The peculiar geographical configuration and position of Italy may indeed increase the exposure of southern regions to exotic pathogens from high-risk areas, such as Northern Africa, Greece, and others. A recent phylogenetic study found indeed that bovine brucellosis isolates from Italy are substantially different than those recovered in northern regions of Europe, and are more closely related to those found in the Middle East (Garofolo et al., 2017). In this study we decided to focus on within-country transmission because of lack of accurate estimates for the risk of introduction of brucellosis from out of the country.

## Conclusion

Bovine brucellosis remains a serious problem in the southern regions of Italy, notwithstanding the long-term and intensive eradication efforts, and causes important economic consequences on the Italian cattle production system, the 2nd largest market in the EU by trade volume. Here we introduced a novel modeling approach assessing the vulnerability of Italian regions to brucellosis persistence, based on empirical movement data and parameterized on outbreak data at farm level. Our findings showed the accuracy of the vulnerability indicator in predicting the number of outbreaks experienced in the affected regions. Most importantly, they identified for the first time animal trade movements as a significant driver for the maintenance of the disease in this area, with an additional appreciable contribution of the use of pastures favoring contiguous spread between neighboring herds. By comparing OBF regions with affected ones, our analysis uncovered the role of limited compliance to trade bans in the southern regions and biosafety measures protecting high production values in the northern regions in explaining the extremely diverse epidemiological situation reported in the country. Providing a synthetic vulnerability indicator, our approach offers a novel tool for veterinary epidemiology based on animal movements and disease spatial transmission that can be readily applied to other epidemiological contexts.

